## Supplementary figures and images for "Viral metagenome characterization reveals species-specific virome profiles in Triatominae populations from the southern United States"

### Supplementary Figure 1

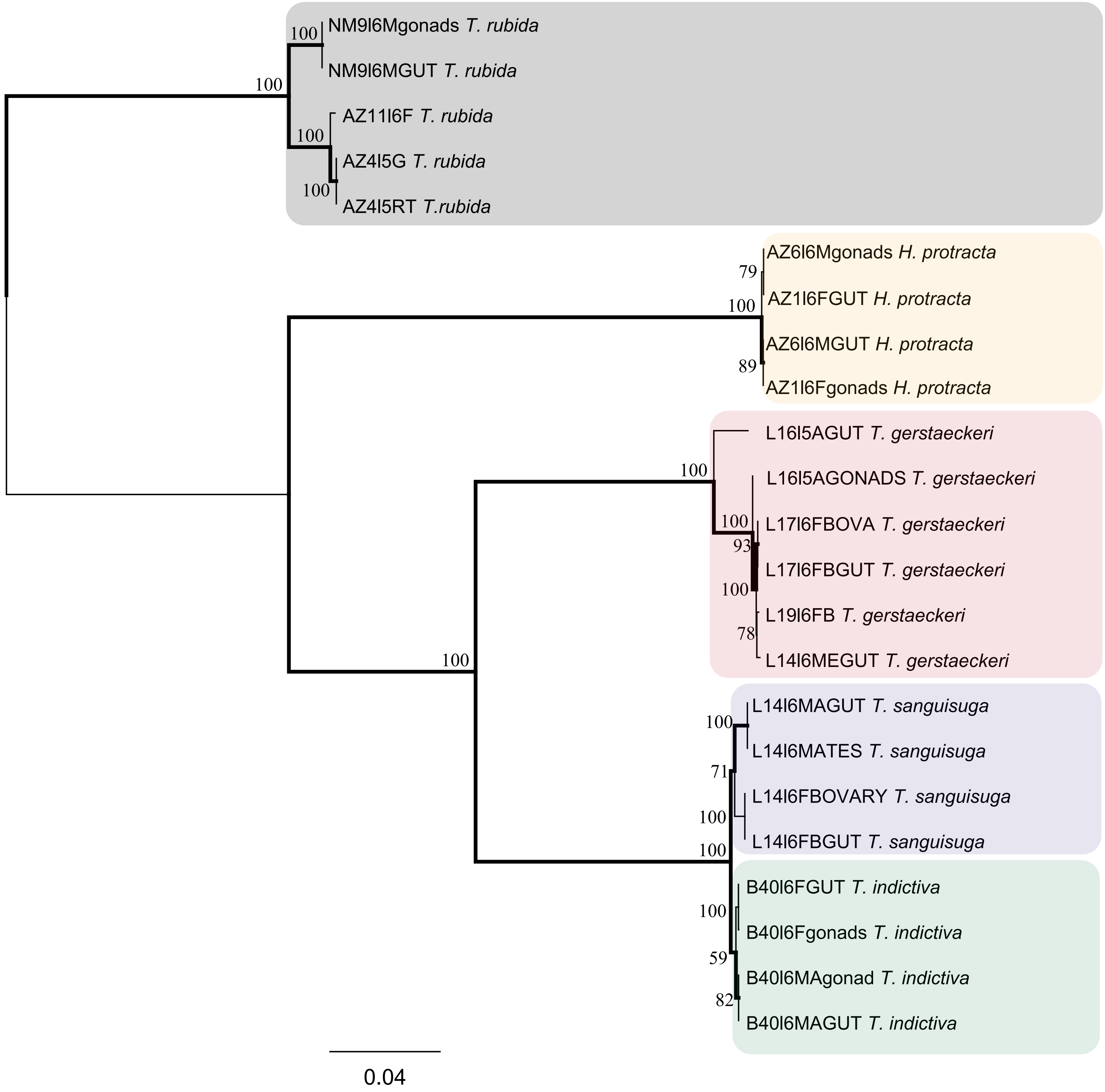
