## Supplementary Figure 2 for "Viral metagenome characterization reveals species-specific virome profiles in Triatominae populations from the southern United States"

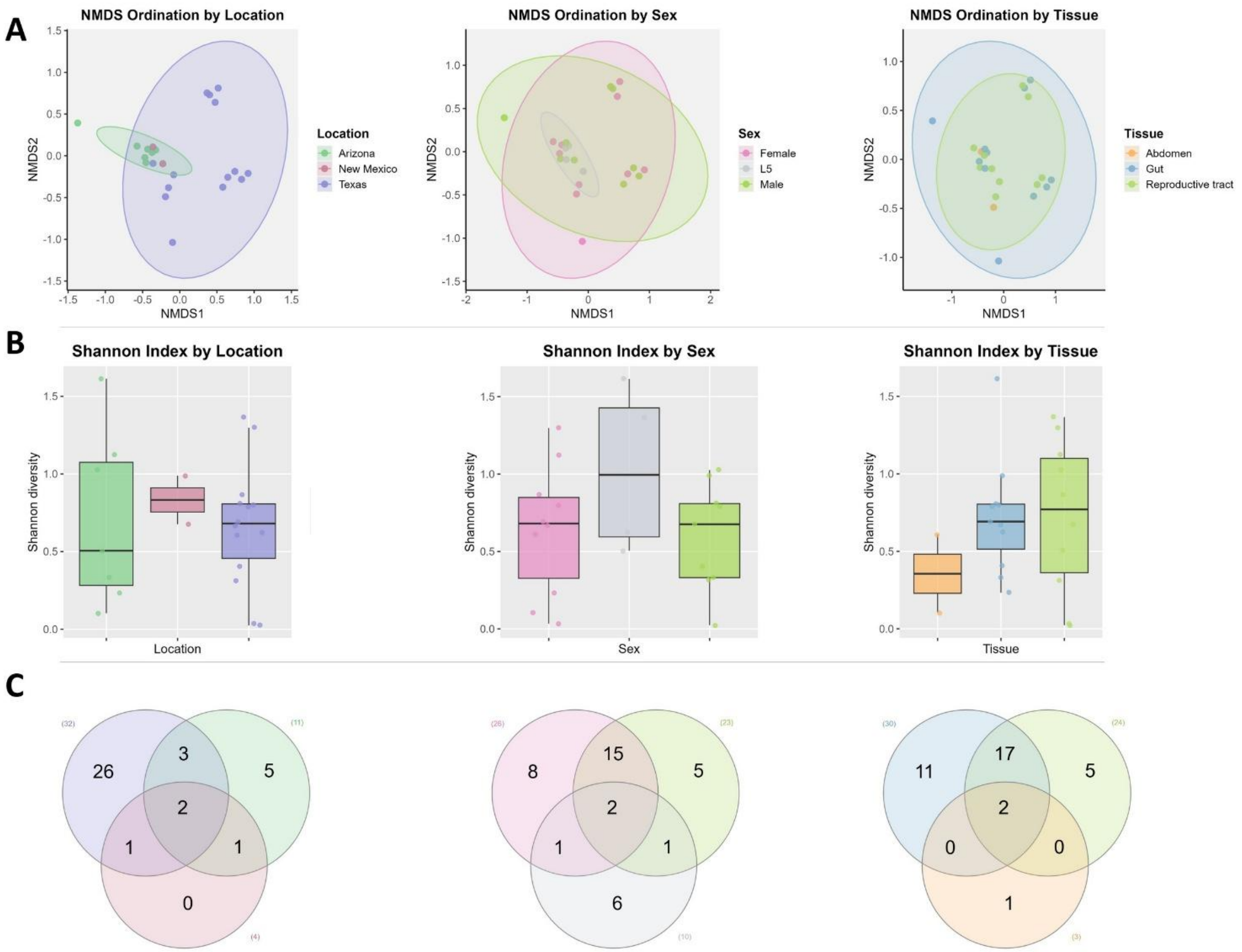


**Supplementary Figure 2. Alpha and beta diversity of vOTUs in natural populations of *Triatoma*. A)** Non-metric multidimensional scaling (NMDS) based on Bray-Curtis distances, showing clustering patterns by study location (PERMANOVA, $p$ = 0.003; $R^2$ = 0.171), sex (PERMANOVA, $p$ = 0.426; $R^2$ = 0.093), and tissue (PERMANOVA, $p$ = 0.97; $R^2$ = 0.138). **B)** Shannon diversity index of vOTUs by location (Kruskal-Wallis test, $p$ = 0.816), sex (Kruskal-Wallis test, $p$ = 0.320), and tissue (Kruskal-Wallis test, $p$ = 0.744). **C)** Venn diagrams displaying the number of shared vOTUs among *Triatoma* populations, grouped by location, sex, and tissue type.
