## Supplementary Table 1 for "Viral metagenome characterization reveals species-specific virome profiles in Triatominae populations from the southern United States"

**Supplementary Table 1. Sample metadata and *T. cruzi* meta-assembly screening.** Summary of 23 *Triatominae* samples with species, sex, geographic location, tissue type, QC, and *T. cruzi* detection.

| Sample ID | Sampling state | Sampling area | Location coordinates | Host species | Stage/Sex | Tissue type | QC | *T. cruzi* meta-assembly screening |
|---|---|---|---|---|---|---|---|---|
| **AZ1I6FGUT** | Arizona | Desert Station | 32.253846N, 111.089947W | *H. protracta* | Female | Gut | PASS | non-infected |
| **AZ1I6Fgonads** | Arizona | Desert Station | 32.253846N, 111.089947W | *H. protracta* | Female | Gonads | PASS | non-infected |
| **AZ6I6MGUT** | Arizona | Desert Station | 32.2540685N, 111.0922009W | *H. protracta* | Male | Gut | PASS | non-infected |
| **AZ6I6Mgonads** | Arizona | Desert Station | 32.2540685N, 111.0922009W | *H. protracta* | Male | Gonads | PASS | non-infected |
| **B40I6FGUT** | Texas | Camp Bullis | 29.7424452N, 98.5876910W | *T. indictiva* | Female | Gut | PASS | non-infected |
| **B40I6Fgonads** | Texas | Camp Bullis | 29.7424452N, 98.5876910W | *T. indictiva* | Female | Gonads | PASS | non-infected |
| **B40I6MAGUT** | Texas | Camp Bullis | 29.7424452N, 98.5876910W | *T. indictiva* | Male | Gut | PASS | non-infected |
| **B40I6MAgonads** | Texas | Camp Bullis | 29.7424452N, 98.5876910W | *T. indictiva* | Male | Gonads | PASS | non-infected |
| **L14I6FBGUT** | Texas | Lackland Air Force Base | 29.3767920N, 98.6849585W | *T. sanguisuga* | Female | Gut | PASS | non-infected |
| **L14I6FBgonads** | Texas | Lackland Air Force Base | 29.3767920N, 98.6849585W | *T. sanguisuga* | Female | Gonads | PASS | non-infected |
| **L14I6MAGUT** | Texas | Lackland Air Force Base | 29.3767920N, 98.6849585W | *T. sanguisuga* | Male | Gut | PASS | non-infected |
| **L14I6Mgonads** | Texas | Lackland Air Force Base | 29.3767920N, 98.6849585W | *T. sanguisuga* | Male | Gonads | PASS | non-infected |
| **L19I6FB** | Texas | Lackland Air Force Base | 29.3675352N, 98.6548306W | *T. gerstaeckeri* | Female | Whole Abdomen | PASS | non-infected |
| **L17I6FBGUT** | Texas | Lackland Air Force Base | 29.3714983N, 98.6571148W | *T. gerstaeckeri* | Female | Gut | PASS | non-infected |
| **L17I6FBgonads** | Texas | Lackland Air Force Base | 29.3714983N, 98.6571148W | *T. gerstaeckeri* | Female | Gonads | PASS | non-infected |
| **L14I6MEGUT** | Texas | Lackland Air Force Base | 29.3767920N, 98.6849585W | *T. gerstaeckeri* | Male | Gut | PASS | non-infected |
| **L16I5AGUT** | Texas | Lackland Air Force Base | 29.3829725N, 98.6891581W | *T. gerstaeckeri* | L5 | Gut | PASS | infected |
| **L16I5Agonads** | Texas | Lackland Air Force Base | 29.3829725N, 98.6891581W | *T. gerstaeckeri* | L5 | Gonads | PASS | non-infected |
| **NM9I6MGUT** | New Mexico | Hatch | 32.6990360N, 107.1797240W | *T. rubida* | Male | Gut | PASS | non-infected |
| **NM9I6Mgonads** | New Mexico | Hatch | 32.6990360N, 107.1797240W | *T. rubida* | Male | Gonads | PASS | non-infected |
| **AZ4I5GUT** | Arizona | Desert Station | 32.2540327N, 111.0917907W | *T. rubida* | L5 | Gut | PASS | non-infected |
| **AZ4I5Rgonads** | Arizona | Desert Station | 32.2540327N, 111.0917907W | *T. rubida* | L5 | Gonads | PASS | non-infected |
| **AZ11I6F** | Arizona | Desert Station | 32.2575005N, 111.0842631W | *T. rubida* | Female | Whole Abdomen | PASS | infected |
