## Supplementary Table 2 for "Viral metagenome characterization reveals species-specific virome profiles in Triatominae populations from the southern United States"

**Supplementary Table 2. Blood meal analysis of *Triatominae* gut tissues.** Twelve DNA isolates from gut samples analysed using vertebrate 12S rRNA gene sequences to identify blood meal sources.

| 12S rRNA sequence | Best blast hit acc.no. | Description | % identity | Length | e-value | note |
|---|---|---|---|---|---|---|
| **NM9i6M_L1085.ab1** | DQ179667.1 | *Neotoma albigula* isolate TK74854 12S ribosomal RNA gene, partial sequence; mitochondrial | 100.000 | 141 | 1.75e-65 | NA |
| **L17i6FB_L1085.ab1** | HM563849.1 | *Incilius signifer* voucher UTA:A-JRM 4968 tRNA-Phe and 12S ribosomal RNA genes, partial sequence; mitochondrial | 100.000 | 133 | 4.67e-61 | equal hits to other Bufonidae, including Incilius nebulifer |
| **L16i5A_L1085.ab1** | JN393214.1 | *Didelphis virginiana* 12S ribosomal RNA gene, partial sequence | 90.071 | 141 | 6.42e-50 | overlapping signal |
| **L14i6ME_L1085.ab1** | ON597633.1 | *Homo sapiens* isolate UniPV_126 mitochondrion, complete genome | 100.000 | 137 | 2.82e-63 | NA |
| **L14i6MA_L1085.ab1** | NA | NA | NA | NA | NA | overlapping signal |
| **L14i6FB_L1085.ab1** | NA | NA | NA | NA | NA | overlapping signal |
| **B40i6MA_L1085.ab1** | MG250549.1 | *Sus scrofa* isolate LUC208 mitochondrion, complete genome | 92.958 | 142 | 1.39e-51 | overlapping signal |
| **B40i6F_L1085.ab1** | ON597633.1 | *Homo sapiens* isolate UniPV_126 mitochondrion, complete genome | 100.000 | 137 | 2.82e-63 | NA |
| **AZ11i6F_L1085.ab1** | DQ179667.1 | *Neotoma albigula* isolate TK74854 12S ribosomal RNA gene, partial sequence; mitochondrial | 100.000 | 141 | 1.75e-65 | NA |
| **AZ6i6M_L1085.ab1** | DQ179667.1 | *Neotoma albigula* isolate TK74854 12S ribosomal RNA gene, partial sequence; mitochondrial | 100.000 | 141 | 1.75e-65 | NA |
| **AZ4i5G_L1085.ab1** | DQ179667.1 | *Neotoma albigula* isolate TK74854 12S ribosomal RNA gene, partial sequence; mitochondrial | 100.000 | 141 | 1.75e-65 | NA |
| **AZ1i6F_L1085.ab1** | DQ179667.1 | *Neotoma albigula* isolate TK74854 12S ribosomal RNA gene, partial sequence; mitochondrial | 100.000 | 141 | 1.75e-65 | NA |
