## Supplementary Table 3 for "Viral metagenome characterization reveals species-specific virome profiles in Triatominae populations from the southern United States"

**Supplementary Table 3. Viral Operational Taxonomic Units (vOTUs).** List of vOTUs detected in Triatominae natural populations from the southern United States.

| vOTU | NCBI acc. number | vOTU abbreviation | Species | length (nt) | ORF length (aa) | Protein domains in the sequence (e-value) * | Assembly level | Closest related sequence | Reported host /country | Virus taxon | Baltimore classification |
|---|---|---|---|---|---|---|---|---|---|---|---|
| **Triatoma sanguisuga chuviridae vOTU1** | PX139058 | Chu_vOTU1 | *T. sanguisuga* | 15597 | 2587 | RNA-directed RNA polymerase (1.2e-158) | complete cds | Guiyang chuvirus 1 (MZ209784.1) | *Cletus punctiger* / China | *Chuviridae* | ssRNA(-) |
| | | | | | 1156 | Fusion glycoprotein (0.0063) | complete cds | | | | |
| | | | | | 914 | RNA virus nucleoprotein (11) | complete cds | | | | |
| **Triatoma sanguisuga chuviridae vOTU2** | PX139059 | Chu_vOTU2 | *T. indictiva* | 10633 | 2218 | RNA-directed RNA polymerase (1.1e-176) | complete cds | Sanya chuvirus 2 (MZ209828.1) | *Nilaparvata lugens* / China | *Chuviridae* | ssRNA(-) |
| | | | | | 471 | Envelope glycoprotein (1.5e-27) | complete cds | | | | |
| | | | | | 462 | RNA virus nucleoprotein (0.027) | complete cds | | | | |
| **Triatoma indictiva chuviridae vOTU3** | PX139060 | Chu_vOTU3 | *T. gersteackeri* | 1542 | 513 | RNA-dependent RNA polymerase (3.7e-37) | partial cds | Neuropteran chu-related virus OKIAV150 (MW288180.1) | *Eumantispa harmandi* / Japan | *Chuviridae* | ssRNA(-) |
| **Triatoma gersteackeri chuviridae vOTU4** | PX139061 | Chu_vOTU4 | *T. gersteackeri* | 1539 | 511 | RNA-dependent RNA polymerase (1.1e-94) | partial cds | Sanya chuvirus 2 (MZ209828.1) | *Nilaparvata lugens* / China | *Chuviridae* | ssRNA(-) |
| **Triatoma gersteackeri chuviridae vOTU5** | PX139062 | Chu_vOTU5 | *T. gersteackeri* | 1505 | 475 | RNA-dependent RNA polymerase (4.7e-20) | partial cds | Sanya chuvirus 2 (MZ209828.1) | *Nilaparvata lugens* / China | *Chuviridae* | ssRNA(-) |
| **Triatoma indictiva chuviridae vOTU6** | PX139063 | Chu_vOTU6 | *T. indictiva* | 1296 | 430 | RNA-dependent RNA polymerase (2.5e-37) | partial cds | Neuropteran chu-related virus OKIAV150 (MW288180.1) | *Eumantispa harmandi* / Japan | *Chuviridae* | ssRNA(-) |
| **Triatoma gersteackeri chuviridae vOTU7** | PX139064 | Chu_vOTU7 | *T. gersteackeri* | 858 | 280 | RNA-dependent RNA polymerase (1.9e-40) | partial cds | Blattodean chu-related virus OKIAV148 (MT153417.1) | *Periplaneta americana*/ Germany | *Chuviridae* | ssRNA(-) |
| **Triatoma gersteackeri virgaviridae vOTU8** | PX139065 | Virga_vOTU8 | *T. gersteackeri* | 9786 | 2933 | Replication proteins (Mtr, Hel and RdRp) (2.7e-66) | complete cds | Xiangshan martelli-like virus 3 (OK491508.1) | Insect / China | *Virgaviridae* | ssRNA(+) |
| | | | | | 220 | Capsid protein (1.2e-31) | complete cds | | | | |
| **Triatoma gersteackeri virgaviridae vOTU9** | PX139066 | Virga_vOTU9 | *T. gersteackeri* | 3682 | 1180 | Polyprotein (Mtr, Hel and RdRp) (4e-41) | partial cds | Xiangshan martelli-like virus 2 (OK491507.1) | Insect / China | *Virgaviridae* | ssRNA(+) |
| **Triatoma gersteackeri virgaviridae vOTU10** | PX139067 | Virga_vOTU10 | *T. gersteackeri* | 2904 | 702 | Polyprotein (Mtr, Hel and RdRp) (3e-67) | partial cds | Xiangshan martelli-like virus 3 (OK491508.1) | Insect / China | *Virgaviridae* | ssRNA(+) |
| | | | | | 213 | Capsid protein (6.9e-32) | complete cds | | | | |
| **Triatoma spp. virgaviridae vOTU11** | PX139068 | Virga_vOTU11 | *T. sanguisuga, T. indictiva* | 1227 | 313 | Genome polyprotein (1.8e-41) | partial cds | Pedersore virga-like virus (ON955134.1) | *Ochlerotatus communis* / Finland | *Virgaviridae* | ssRNA(+) |
| | | | | | 74 | Capsid protein (53) | complete cds | | | | |
| **Triatoma spp. virgaviridae vOTU12** | PX139069 | Virga_vOTU12 | *T. sanguisuga, T. indictiva* | 1161 | 214 | RNA-dependent RNA polymerase (1.7e-31) | partial cds | Ginka virga-like virus (ON860475.1) | *Ochlerotatus communis* / Sweden | *Virgaviridae* | ssRNA(+) |
| **Triatoma spp. elliovirales vOTU13R** | PX139070/ PX139071 | Ellio_vOTU13 | *T. sanguisuga, T. gersteackeri* | 7919 | 2577 | Bunyavirus RNA dependent RNA polymerase (4.81e-20) | complete cds | Orius laevigatus bunyavirus 2 (PP908622.1) | *Orius laevigatus* / Spain | Elliovirales | ssRNA(-) |
| | | | | 4670 | 1065 | Envelopment polyprotein (2.5e-73) | complete cds | | | | |
| **Triatoma spp. elliovirales vOTU14** | PX139072/ PX139073 | Ellio_vOTU14 | *T. sanguisuga, T. gersteackeri* | 7908 | 2569 | Bunyavirus RNA dependent RNA polymerase (1.41e-20) | complete cds | Orius laevigatus bunyavirus 2 (PP908622.1) | *Orius laevigatus* / Spain | Elliovirales | ssRNA(-) |
| | | | | 3499 | 1058 | Glycoprotein (5.8e-79) | partial cds | | | | |
| **Hospesneotomae protracta arenaviridae vOTU15** | PX139074/ PX139075 | Arena_vOTU15 | *H. protracta* | 7182 | 2220 | Arenavirus RNA polymerase (0e+00), Arenavirus cap snatching domain (3.33e-57) | complete cds | Big brushy tank virus (EU938666.1) | *Neotoma albigula* / USA | *Arenaviridae* | ssRNA(-) |
| | | | | | 95 | RING finger protein Z (5.9e-53) | complete cds | | | | |
| | | | | 3409 | 576 | Nucleoprotein (8.6e-208), Glycoprotein (4e-148) | partial cds | | | | |
| **Triatoma rubida arenaviridae vOTU16** | PX139076/ PX139077 | Arena_vOTU16 | *T. rubida* | 1261 | 236/ 181 | RNA-directed RNA polymerase (4.9e-56) | partial cds | Mammarenavirus whitewaterense (NC_010703.1) | *Neotoma albigula* / USA | *Arenaviridae* | ssRNA(-) |

| | | | | | | | | | | | |
|---|---|---|---|---|---|---|---|---|---|---|---|
| **Triatoma indictiva astroviridae vOTU17** | PX139078 | Astro_vOTU17 | *T. indictiva* | 5904 | 662 | Viral protease (0.00075) | partial cds | Flumine Astrovirus 3 (OM954094.1) | River / New Zealand | *Astroviridae* | ssRNA(+) |
| | | | | | 546 | RNA-dependent RNA polymerase (2.9e-50) | complete cds | | | | |
| | | | | | 389 | Capsid protein (1.2) | complete cds | | | | |
| **Hospesneotomae protracta benyviridae vOTU18** | PX139079 | Beny_vOTU18 | *H. protracta* | 5340 | 1387 | Polyprotein (Mtr, Hel and RdRp) (8e-32) | complete cds | Sanya benyvirus 1 (MZ209861.1) | *Sesamia inferens*/ China | *Benyviridae* | ssRNA(+) |
| | | | | | 244 | Virus coat protein (4e-30) | complete cds | | | | |
| **Triatoma gersteackeri benyviridae vOTU19** | PX139080 | Beny_vOTU19 | *T. gersteackeri* | 5297 | 1384 | Polyprotein (Mtr, Hel and RdRp) (3.6e-47) | complete cds | Guiyang benyvirus 1 (MZ209815.1) | *Harmonia axyridis*/ China | *Benyviridae* | ssRNA(+) |
| | | | | | 253 | Virus coat protein (1e-30) | complete cds | | | | |
| **Triatoma gersteackeri benyviridae vOTU20** | PX139081 | Beny_vOTU20 | *T. gersteackeri* | 5280 | 1384 | Polyprotein (Mtr, Hel and RdRp) (4.2e-48) | complete cds | Benyviridae sp. (PQ792268.1) | *Bradysia coprophila*/ China | *Benyviridae* | ssRNA(+) |
| | | | | | 253 | Virus coat protein (1.2e-30) | complete cds | | | | |
| **Hospesneotomae protracta benyviridae vOTU21** | PX139082 | Beny_vOTU21 | *H. protracta* | 2290 | 667 | Replicase polyprotein (Mtr, Hel and RdRp) (7.8e-32) | partial cds | Benyviridae sp. (PQ792268.1) | *Bradysia coprophila*/ China | *Benyviridae* | ssRNA(+) |
| **Triatoma spp. tombusviridae vOTU22** | PX139083 | Tombus_vOTU22 | *T. sanguisuga, T. indictiva* | 3612 | 505 | RNA-dependent RNA polymerase (1.4e-41) | complete cds | Hemipteran tombus-related virus (QTJ63611.1) | *Notostira elongata* / Germany | *Tombusviridae* | ssRNA(+) |
| | | | | | 186 | Coat protein (1e-31) | complete cds | | | | |
| **Triatoma indictiva narnaviridae vOTU23** | PX139084 | Narna_vOTU23 | *T. indictiva* | 2993 | 921 | RNA-directed RNA polymerase (2.6e-21) | complete cds | Wuhan spider virus 7 (NC_033702.1) | Spiders / China | *Narnaviridae* | ssRNA(+) |
| **Triatoma indictiva narnaviridae vOTU24** | PX139085 | Narna_vOTU24 | *T. indictiva* | 2705 | 824 | RNA-dependent RNA polymerase (5.5e-20) | complete cds | Serbia narna-like virus 3 (MT822185.1) | *Culex pipiens* / Serbia | *Narnaviridae* | ssRNA(+) |
| **Triatoma indictiva solemoviridae vOTU25** | PX139086 | Solemo_vOTU25 | *T. indictiva* | 2936 | 501 | Protease (6.6e-13) | complete cds | Atrato Sobemo-like virus 2 (MN661089.1) | *Culex* sp / Colombia | *Solemoviridae* | ssRNA(+) |
| | | | | | 490 | RNA dependent RNA polymerase (7.6e-46) | complete cds | | | | |
| **Triatoma sanguisuga solemoviridae vOTU26** | PX139087 | Solemo_vOTU26 | *T. sanguisuga* | 2933 | 562 | Peptidase (6.9e-15) | complete cds | Rhodnius prolixus virus 6 (MZ328309.1) | *Rhodnius prolixus* / Brazil | *Solemoviridae* | ssRNA(+) |
| | | | | | 506 | RNA-dependent RNA polymerase (2.6e-39) | complete cds | | | | |
| **Triatoma indictiva rhabdoviridae vOTU27** | PX139088 | Rhabdo_vOTU27 | *T. indictiva* | 1856 | 617 | RNA dependent RNA polymerase (5e-108) | partial cds | Sanya conocephalus maculatus rhabdovirus 1 (MZ209844.1) | *Conocephalus maculatus* / China | *Rhabdoviridae* | ssRNA(-) |
| **Triatoma spp. rhabdoviridae vOTU28** | PX139089 | Rhabdo_vOTU28 | *T. sanguisuga, T. indictiva* | 1273 | 422 | RNA-dependent RNA polymerase (1e-25) | partial cds | Blattodean rhabdo-related virus OKIAV14 (MT153532.1) | *Deropeltis erythrocephala*/ Germany | *Rhabdoviridae* | ssRNA(-) |
| **Triatoma indictiva partitiviridae vOTU29** | PX139090 | Partiti_vOTU29 | *T. indictiva* | 1800 | 562 | RNA-dependent RNA polymerase (1.9e-36) | partial cds | Hubei partiti-like virus 10 (KX884114.1) | Odonata / China | *Partitiviridae* | dsRNA |
| **Triatoma spp. partitiviridae vOTU30** | PX139091 | Partiti_vOTU30 | *T. sanguisuga, T. indictiva* | 997 | 252 | RNA-dependent RNA polymerase (7.09e-27) | partial cds | Hangzhou partitivirus 1 (MZ209755.1) | *Orthetrum chrysostigma* / China | *Partitiviridae* | dsRNA |
| **Triatoma sanguisuga partitiviridae vOTU31** | PX139092 | Partiti_vOTU31 | *T. sanguisuga* | 590 | 171 | RNA-dependent RNA polymerase (7.9e-7) | partial cds | Hubei partiti-like virus 10 (KX884114.1) | Odonata / China | *Partitiviridae* | dsRNA |
| **Triatominae partitiviridae vOTU32** | PX139093 | Partiti_vOTU32 | *H. protracta, T. sanguisuga, T. indictiva* | 522 | 167 | RNA dependent RNA polymerase (4.7e-19) | partial cds | Hubei partiti-like virus 11 (NC_032139.1) | Odonata / China | *Partitiviridae* | dsRNA |
| **Triatoma indictiva circoviridae vOTU33** | PX139094 | Circo_vOTU33 | *T. indictiva* | 998 | 331 | Viral coat protein (S domain) (1.27e-08) | partial cds | Circovirus sp. (OP564891.1) | *Neophema chrysogaster* / Australia | *Circoviridae* | ssDNA |
| **Triatoma spp. circoviridae vOTU34** | PX139095 | Circo_vOTU34 | *T. rubida, T. sanguisuga, T. indictiva* | 992 | 167 | Viral coat protein (S domain) (2.9e-17) | partial cds | Circovirus sp. (OP564891.1) | water / New Zealand | *Circoviridae* | ssDNA |
| **Triatoma spp. circoviridae vOTU35** | PX139096 | Circo_vOTU35 | *T. rubida, T. sanguisuga* | 568 | 188 | Rep protein (2e-23) | partial cds | Circovirus sp. gt3aAU ( PQ754361.1) | *Ixodes ovatus*/ China | *Circoviridae* | ssDNA |
| **Triatoma spp. circoviridae vOTU36** | PX139097 | Circo_vOTU36 | *T. rubida, T. indictiva* | 507 | 97 | viral replication protein (2.02e-13) | partial cds | Cressdnaviricota sp. (MH616905.1) | red snapper / USA | *Circoviridae* | ssDNA |

| | | | | | | | | | | | |
|---|---|---|---|---|---|---|---|---|---|---|---|
| **Triatoma rubida microviridae vOTU37** | PX139098 | Micro_vOTU37 | *T. rubida* | 734 | 223 | Capsid protein VP1 (3.1e-53) | partial cds | Microvirus sp. (OR349802.1) | *Potamopyrgus antipodarum* / New_Zealand | *Microviridae* | ssDNA |
| **Triatominae orthomyxoviridae vOTU38** | PX139099 | Ortho_vOTU38 | *H. protracta, T. indictiva* | 516 | 171 | Polymerase PB2 (7.9e-36) | partial cds | Halyomorpha halys orthomyxo-like virus 1 (URQ09140.1) | *Halyomorpha halys*/ Italy | *Orthomyxoviridae* | ssRNA(-) |
| **Hospesneotomae protracta orthomyxoviridae vOTU39** | PX139100 | Ortho_vOTU39 | *H. protracta* | 354 | 117 | Nucleoprotein (2.2e-17) | partial cds | Hemipteran orthomyxo-related virus OKIAV191 (QMP82198.1) | *Acanthosoma haemorrhoidale*/ Germany | *Orthomyxoviridae* | ssRNA(-) |
| **Triatoma spp. xinmoviridae vOTU40** | PX139101 | Xinmo_vOTU40 | *T. rubida, T. gersteackeri* | 514 | 137 | RNA dependent RNA polymerase (2e-19) | partial cds | Hangzhou cletus punctiger xinmovirus 1 (UHK03158.1) | *Cletus punctiger*/ China | *Xinmoviridae* | ssRNA(-) |
| **Triatoma rubida rhabdoviridae vOTU41** | PX139102 | Rhabdo_vOTU41 | *T. rubida* | | 278 | Mononegavirales RNA dependent RNA polymerase (4e-26) | partial cds | Blattodean rhabdo-related virus OKIAV14 (QMP82336.1) | *Deropeltis erythrocephala*/ Germany | *Rhabdoviridae* | ssRNA(-) |
