## Supplementary Table 4 for "Viral metagenome characterization reveals species-specific virome profiles in Triatominae populations from the southern United States"

**Supplementary Table 4. Sequencing metrics.** Summary of raw sequencing reads, reads retained after quality control, total number of assembled contigs, and number of viral reads per sample.

| Sample name | Raw reads | Reads after QC | Percentage of reads recovered | Assembled contigs per sample | Viral reads per sample | Viral reads % |
|---|---|---|---|---|---|---|
| **AZ4I5G** | 52525874 | 42241734 | 80.42081127 | 205948 | 3845 | 5.51E-05 |
| | 52525874 | 42372674 | 80.67009794 | | | |
| **AZ4I5RT** | 51414803 | 45067671 | 87.65504946 | 330712 | 80844 | 0.000765962 |
| | 51414803 | 45067671 | 87.65504946 | | | |
| **L16I5AGONADS** | 52705047 | 46110738 | 87.48827793 | 350410 | 545173 | 0.006385778 |
| | 52705047 | 47261468 | 89.67161722 | | | |
| **L16I5AGUT** | 51222221 | 43542511 | 85.00707339 | 432263 | 61593 | 0.000837012 |
| | 51222221 | 43542511 | 85.00707339 | | | |
| **NM9I6MGUT** | 54734683 | 44659157 | 81.59206293 | 199336 | 13213 | 1.514607526 |
| | 54734683 | 44659157 | 81.59206293 | | | |
| **NM9I6Mgonads** | 54631017 | 35627128 | 65.21410356 | 397764 | 27462 | 0.000444255 |
| | 54631017 | 35627128 | 65.21410356 | | | |
| **L14I6FBGUT** | 50778094 | 41898787 | 82.51350868 | 230909 | 4264116 | 0.056845617 |
| | 50778094 | 41898787 | 82.51350868 | | | |
| **L14I6FBOVARY** | 59722382 | 48629329 | 81.42563537 | 344303 | 17590 | 0.000203167 |
| | 59722382 | 48629329 | 81.42563537 | | | |
| **L14I6MAGUT** | 52282492 | 47160673 | 90.20356757 | 266420 | 161984915 | 1.897688314 |
| | 52282492 | 47160673 | 90.20356757 | | | |
| **L14I6MATES** | 51084931 | 37190250 | 72.80082261 | 288631 | 2273 | 3.01E-05 |
| | 51084931 | 37190250 | 72.80082261 | | | |
| **AZ6I6MGUT** | 51069430 | 47833025 | 93.66273522 | 304677 | 1323549 | 0.015621404 |
| | 51069430 | 47833025 | 93.66273522 | | | |
| **AZ6I6Mgonads** | 60944909 | 30463498 | 49.98530394 | 261758 | 8951 | 0.000151405 |
| | 60944909 | 30463498 | 49.98530394 | | | |
| **AZ1I6FGUT** | 55677990 | 47688530 | 85.6505955 | 226184 | 3249 | 4.00E-05 |
| | 55677990 | 47688530 | 85.6505955 | | | |
| **AZ1I6Fgonads** | 52346468 | 43801616 | 83.67635425 | 230695 | 968892 | 0.012557295 |
| | 52346468 | 43801616 | 83.67635425 | | | |

| | | | | | | |
|---|---|---|---|---|---|---|
| **L17I6FBGUT** | 48325959 | 43748312 | 90.52756097 | 490561 | 103194 | 0.001295439 |
| | 48325959 | 43748312 | 90.52756097 | | | |
| **L17I6FBOVA** | 49590199 | 30471507 | 61.44663182 | 268386 | 138320 | 2.59100499 |
| | 49590199 | 30471507 | 61.44663182 | | | |
| **B40I6MAGUT** | 51837806 | 44230870 | 85.32550548 | 191623 | 444998939 | 5.515520683 |
| | 51837806 | 44230870 | 85.32550548 | | | |
| **B40I6MAgonad** | 49369024 | 28137323 | 56.99388143 | 216727 | 5277 | 9.30E-05 |
| | 49369024 | 28137323 | 56.99388143 | | | |
| **B40I6FGUT** | 45515153 | 36184792 | 79.50053908 | 163413 | 298760 | 0.00454642 |
| | 45515153 | 36184792 | 79.50053908 | | | |
| **B40I6Fgonads** | 47414182 | 41034885 | 86.54559305 | 235945 | 17175 | 0.000231871 |
| | 47414182 | 41034885 | 86.54559305 | | | |
| **L14I6MEGUT** | 51437951 | 45934896 | 89.30156646 | 531140 | 33632360 | 0.39191126 |
| | 51437951 | 45934896 | 89.30156646 | | | |
| **AZ11I6F** | 47693246 | 31041027 | 65.08474386 | 394736 | 102274 | 0.002222757 |
| | 47693246 | 31041027 | 65.08474386 | | | |
| **L19I6FB** | 36279160 | 21497807 | 59.25662832 | 186092 | 2100944 | 0.069029902 |
| | 36279160 | 21497807 | 59.25662832 | | | |
