## Supplementary Table 5 for "Viral metagenome characterization reveals species-specific virome profiles in Triatominae populations from the southern United States"

**Supplementary Table 5**. SIMPER analysis. Contribution of each Viral Operational Taxonomic Units (vOTUs) to the Bray-Curtis dissimilarities observed in *Triatoma* natural populations between species (*H.protracta* , *T.rubida, T. indictiva, T.sanguisuga, T.gersteackeri*).

| vOTU | Taxon | Av. dissim | Contrib. % | Cumulative % |
|---|---|---|---|---|
| Solemo_vOTU25 | *Solemoviridae* | 26.62 | 26.8 | 26.8 |
| Solemo_vOTU26 | *Solemoviridae* | 14.64 | 14.73 | 41.54 |
| Ellio_vOTU13 | *Elliovirales* | 10.61 | 10.68 | 52.22 |
| Ellio_vOTU14 | *Elliovirales* | 7.836 | 7.888 | 60.1 |
| Rhabdo_vOTU41 | *Rhabdoviridae* | 7.79 | 7.842 | 67.95 |
| Narna_vOTU24 | *Narnaviridae* | 6.302 | 6.344 | 74.29 |
| Arena_vOTU15 | *Arenaviridae* | 4.64 | 4.671 | 78.96 |
| Beny_vOTU19 | *Benyviridae* | 3.019 | 3.039 | 82 |
| Xinmo_vOTU40 | *Xinmoviridae* | 2.332 | 2.347 | 84.35 |
| Virga_vOTU8 | *Virgaviridae* | 2.066 | 2.08 | 86.43 |
| Chu_vOTU3 | *Chuviridae* | 1.6 | 1.611 | 88.04 |
| Ortho_vOTU38 | *Orthomyxoviridae* | 1.213 | 1.221 | 89.26 |
| Chu_vOTU4 | *Chuviridae* | 1.143 | 1.15 | 90.41 |
| Ortho_vOTU39 | *Orthomyxoviridae* | 1.068 | 1.075 | 91.48 |
| Virga_vOTU10 | *Virgaviridae* | 1.054 | 1.061 | 92.54 |
| Chu_vOTU7 | *Chuviridae* | 1.05 | 1.057 | 93.6 |
| Chu_vOTU5 | *Chuviridae* | 1.033 | 1.04 | 94.64 |
| Beny_vOTU18 | *Benyviridae* | 0.9424 | 0.9487 | 95.59 |
| Virga_vOTU9 | *Xinmoviridae* | 0.9375 | 0.9438 | 96.53 |
| Circo_vOTU35 | *Circoviridae* | 0.7596 | 0.7647 | 97.3 |
| Arena_vOTU16 | *Arenaviridae* | 0.5447 | 0.5484 | 97.85 |
| Partiti_vOTU32 | *Partitiviridae* | 0.3718 | 0.3743 | 98.22 |
| Circo_vOTU34 | *Circoviridae* | 0.341 | 0.3433 | 98.56 |
| Micro_vOTU37 | *Microviridae* | 0.2801 | 0.282 | 98.85 |
| Chu_vOTU2 | *Chuviridae* | 0.2587 | 0.2605 | 99.11 |
| Virga_vOTU12 | *Virgaviridae* | 0.1995 | 0.2009 | 99.31 |
| Circo_vOTU36 | *Circoviridae* | 0.1522 | 0.1532 | 99.46 |
| Virga_vOTU11 | *Virgaviridae* | 0.1185 | 0.1193 | 99.58 |
| Tombus_vOTU22 | *Tombusviridae* | 0.1089 | 0.1097 | 99.69 |
| Partiti_vOTU31 | *Partitiviridae* | 0.07963 | 0.08016 | 99.77 |
| Beny_vOTU20 | *Benyviridae* | 0.07289 | 0.07338 | 99.84 |
| Astro_vOTU17 | *Astroviridae* | 0.06069 | 0.06109 | 99.9 |
| Beny_vOTU21 | *Benyviridae* | 0.03512 | 0.03535 | 99.94 |
| Partiti_vOTU30 | *Partitiviridae* | 0.03346 | 0.03368 | 99.97 |
| Narna_vOTU23 | *Narnaviridae* | 0.009685 | 0.00975 | 99.98 |
| Rhabdo_vOTU29 | *Rhabdoviridae* | 0.007088 | 0.007136 | 99.99 |
| Chu_vOTU1 | *Chuviridae* | 0.005412 | 0.005448 | 100 |
| Chu_vOTU6 | *Circoviridae* | 0.002119 | 0.002133 | 100 |
| Circo_vOTU33 | *Circoviridae* | 0.001271 | 0.00128 | 100 |
| Rhabdo_vOTU27 | *Rhabdoviridae* | 0.000618 | 0.0006222 | 100 |
| Rhabdo_vOTU28 | *Rhabdoviridae* | 0.000252 | 0.0002537 | 100 |
